# The L1 Cell Adhesion Molecule Constrains Dendritic Spine Density through Ankyrin Binding in Pyramidal Neurons of the Mouse Cerebral Cortex

**DOI:** 10.1101/2022.05.16.492130

**Authors:** Kelsey E. Murphy, Sarah D. Wade, Justin E. Sperringer, Vishwa Mohan, Bryce W. Duncan, Yubin Pak, David Lutz, Melitta Schachner, Patricia F. Maness

## Abstract

A novel function for L1 cell adhesion molecule and its interaction with Ankyrin, an actin-spectrin adaptor protein, was identified in constraining dendritic spine density on pyramidal neurons in the mouse neocortex. In an L1-null mouse mutant increased spine density was observed on apical but not basal dendrites of pyramidal neurons in diverse cortical areas (prefrontal cortex layer 2/3, motor cortex layer 5, visual cortex layer 4).The Ankyrin binding motif (FIGQY) in L1’s cytoplasmic domain was critical for spine formation, as demonstrated by increased spine density in the prefrontal cortex of a mouse mutant (L1YH) harboring a tyrosine to histidine mutation in this motif, which disrupts L1-Ankyrin association. This mutation is a known variant in the human L1 syndrome. In both mutants mature mushroom spines rather than immature spines were predominant. L1 was detected in spines and dendrites of wild-type prefrontal cortical neurons by immmunostaining. L1 coimmunoprecipitated with Ankyrin B (220 kDa) from cortical lysates of wild-type but not L1YH mice. Spine pruning assays in cortical neuron cultures from wild-type and L1YH mutant mice showed that the L1-Ankyrin interaction mediated spine retraction in response to the class 3 Semaphorins, Sema3F and to a lesser extent Sema3B. These ligands also induce spine pruning through other L1 family adhesion molecules, NrCAM and Close Homolog of L1 (CHL1), respectively. This study provides insight into the molecular mechanism of spine regulation and underscore the potential for this adhesion molecule to regulate cognitive and other L1-related functions that are abnormal in the L1 syndrome.

## Introduction

Dendritic spines on cortical pyramidal neurons receive 80-90% of excitatory glutamatergic synapses in the neocortex (1). Dendritic spine number is tightly regulated in developing and adult brain to achieve an appropriate balance of excitatory and inhibitory connections that are essential for cortical functioning. Patients with autism spectrum disorder (ASD) or Fragile X Syndrome display an elevated spine density in the prefrontal cortex (PFC), where essential circuits contribute to social behavior and cognition (2-6), while in the PFC of patients with schizophrenia spine density is reduced (7). Decreased spine density and atypical spine morphology have also been described in subjects with cognitive impairment (8) and Down’s syndrome (reviewed in (9)). During development of the human and mouse brain, spines are initially overproduced, eliminated in substantial numbers during adolescence, and stabilized in adulthood (10-13). An attractive hypothesis is that aberrant spine pruning and/or growth during adolescence leads to neuropsychiatric deficits. Therefore defining the molecules and mechanisms regulating dendritic spines in neocortical circuits is important for understanding normal maturation and pathological consequences of deleterious mutations.

L1 family cell adhesion molecules (L1, Close Homolog of L1 (CHL1), Neuron-glial related CAM (NrCAM), and Neurofascin) are transmembrane recognition molecules that perform diverse functions in neural development including axon growth and guidance, neuronal migration, cell survival, and synaptic plasticity (14, 15). Mutations in the L1 gene on the human X chromosome are linked to a syndrome of severe intellectual disability accompanied by hydrocephalus, aphasia, and spastic paraplegia with an incidence of 1/25,000-1/60,000 males (16, 17). Over 250 distinct L1 syndrome mutations have been identified in all regions of the gene, most of which result in loss of function (18, 19). L1 null mutant mice as a model exhibit L1 syndrome-related features including axonal misguidance and enlarged brain ventricles (20-23). One human pathological mutation in the L1 syndrome results in a tyrosine to histidine substitution in a cytoplasmic motif (FIGQY^1229^) that is highly conserved among L1 family members (24). The L1 FIGQY motif mediates reversible binding of the actin-spectrin adaptor Ankyrin (25-27). Ankyrin B is encoded by the *Ank2* gene, a high confidence ASD gene (Simons Foundation SFARI database). An L1 knock-in mouse harboring the FIGQH mutation displays axon targeting errors (28) and impaired stabilization of interneuron synapses (29-31).

Recent studies with L1-CAM-deficient mouse models have revealed novel roles for NrCAM and CHL1 in constraining the density of dendritic spines and excitatory synapses on apical dendrites of cortical pyramidal neurons in the PFC (32-35). Distinct spine subpopulations are pruned in response to secreted Semaphorin-3 ligands Sema3F and Sema3B through receptor complexes comprising NrCAM/Neuropilin2/PlexinA3 and CHL1/Neuropilin2/PlexinA4, respectively. A potential function for L1 in regulating spine density has not been examined. To investigate a role for L1 and its interaction with Ankyrin in dendritic spine regulation, we analyzed L1-null and L1YH mice for alterations in spine density, morphology, and dendritic arborization in cortical pyramidal neurons. We found that deletion of L1 or mutation of the Ankyrin binding site on L1 increased the density of spines on apical, but not basal dendrites of pyramidal neurons in the PFC and other neocortical areas.

## Materials and Methods

### Mice

L1-deficient knockout mice (20) were bred hemizygously on the SV129 genetic background and housed at 22°C on a 12 hr light/dark cycle with *ad libitum* access to food and water. The L1 gene is on the X chromosome, thus hemizygous males (L1-/y) are null mutants. L1-/y and wild type (WT) male littermates were analyzed in this study. L1Y^1229^H (L1YH) mice, mutated in the Ankyrin binding motif FIGQY (28), were maintained by crossing WT C57Bl/6 males to heterozygous L1YH females (C57Bl/6) to yield WT and mutant littermate males, because L1-/y males have greatly reduced fertility. WT and L1 mutant mice were analyzed in adulthood (postnatal days P50-150) after the most active period of juvenile spine remodeling has been completed (10, 36).

For immunostaining Nex1Cre-ERT2:RCE mice containing a loxP-stop-loxP EGFP allele were induced to express EGFP in pyramidal neurons by daily intraperitoneal injections of tamoxifen (100 mg/kg) at P10-P13 as described (37). Postnatal tamoxifen induction in Nex1-CreERT2:RCE mice has been shown to achieve cell-specific targeting of postmitotic cortical pyramidal neurons with no detectable targeting of interneurons, oligodendroglia, astrocytes, or non-neural cells (37). All mice were handled according to the University of North Carolina Institutional Animal Care and Use Committee policies in accordance with NIH guidelines.

### Golgi Impregnation and Spine Analysis

Adult mice (50-150) were anesthetized with isoflurane, brains were isolated and rinsed in distilled water, then processed for Golgi impregnation using the FD Rapid Golgi Stain Kit (FD NeuroTechnologies) as described (34). Coronal vibratome sections (100 µm) containing the medial PFC (cingulate cortex 1 and 2), M1, and V1 were mounted on gelatin-coated microscope slides. Golgi-labeled neurons were imaged under brightfield illumination using an Olympus Neville microscope by scanning optical sections at 60x and generating minimum intensity projections in FIJI. Spine density and arborization of pyramidal cell dendrites were quantified using Neurolucida software (MBF Bioscience) by investigators blind to genotype as described (32-34). Briefly, spines were traced and quantified on 30 µm segments of the first branch of apical or basal dendrites from confocal z-stack images (6-8 mice/genotype; 30-55 neurons/genotype; 10-30 spines/neuron). Mean spine number per 10 µm of dendritic length (density) was calculated. Mean spine densities/10 μm ± SEM were compared by Mann-Whitney two-tailed t-tests (unequal variance, p<0.05), as normal distributions were not assumed. Different spine morphologies are classified as mushroom, stubby, and thin spines based on the relative size of the spine head and neck (38, 39). The density of spines of each morphological type was scored blind to observer using Neurolucida software on apical dendrite segments from images of Golgi-labeled pyramidal neurons of WT, L1-/y, and L1YH mice at P50 as defined (38, 39) and described previously (32). The densities of each spine type were calculated and compared for significant differences by two-tailed Mann-Whitney t-tests (p < 0.05).

Dendritic arborization of Golgi-labeled neurons was assessed by Sholl analysis of image stacks captured at lower magnification (20x/0.5 NA, 1 µm z-series sections). The Sholl center was defined as the midpoint of the cell body at a soma detector sensitivity of 1.5 µm, and the automatic tracing mode was used to seed and trace dendritic arbors. Images in DAT format were subjected to Sholl analysis using Neurolucida Explorer with a starting radius of 10 µm and radius increments of 10 µm. Two factor ANOVA was used to assess significant differences in the mean number of crossings at each distance from the soma with significance set at p < 0.05.

### Immunofluorescence Staining

Nex1Cre: RCE mice (WT) were induced with tamoxifen at P10-P13 and brains were harvested at P20. After transcardial perfusion with 4% PFA, fixed brains were isolated, sectioned, permeabilized, and blocked in 10% normal donkey serum containing 0.3% Triton X-100 in PBS. Sections were stained with antibodies directed against GFP (1:250, Abcam #13970, RRID:AB_300798, host chicken), L1 (1:100, monoclonal antibody 324, Millipore-Sigma #MAB5272, RRID:AB_2133200, host rat) and MAP2 (1:100, Abcam 254143, host mouse) for 2 hours at 4°C. AlexaFluor-conjugated secondary antibodies (anti-chicken AF488, anti-rat AF555, anti-mouse AF647 (1:250, Thermofisher) were incubated with sections for 2 hours at room temperature. Sections were mounted on Superfrost Plus slides with Prolong Glass mountant, cured for 48-72 hours, imaged confocally and deconvolved using AutoQuant 3 software (Media Cybernetics) with default deconvolution settings in Imaris (Bitplane). Microscopy was performed in the UNC Microscopy Services Laboratory (Dr. Pablo Ariel, Director).

### Cortical lysates, Immunoprecipitation, and Immunoblotting

For preparation of mouse cortical lysates, forebrains (P30) were homogenized in RIPA buffer, incubated for 15 minutes on ice, and centrifuged at 16,000 x g for 10 minutes. The supernatant was retained, and protein concentration was determined by BCA. For immunoprecipitation, lysates (1 mg) were precleared for 30 minutes at 4°C using Protein A/G Sepharose beads (ThermoFisher). Precleared lysates (equal amounts of protein) were incubated with 2.5 µg nonimmune IgG (NIgG) or L1 monoclonal antibody 2C2 (Abcam #ab24345) together with 1.25 µg L1 monoclonal antibody 5G3 (BD/Pharmingen #554273) for 2 hours on ice. Protein A/G Sepharose beads were added for 30 minutes before washing with RIPA buffer. Samples were subjected to SDS-PAGE (6%) and transferred to nitrocellulose. Membranes were blocked in TBST containing 5% nonfat dried milk and incubated overnight with monoclonal antibodies (1:1000) against Ankyrin B (ThermoFisher #33-3700, RRID:AB_2533115), washed, and incubated with HRP-secondary antibodies (1:5000) for 1 h. Blots were developed using Western Bright ECL Substrate (Advansta) and exposed to film for times yielding a linear response of signal.

### Cortical Neuron Cultures and Spine Retraction Assay

Cortical neuron cultures were prepared from forebrains of WT and L1YH embryos at E15.5 and as described (32, 33). At DIV11 cells were transfected with pCAG-IRES-EGFP to aid in visualizing spines of neuronal cells (32, 35). At DIV 14 cultures were treated with purified human Fc or recombinant mouse Sema3F-Fc or Sema3B-Fc fusion proteins (R&D Systems) at 5 nM for 30 minutes. Cultures were fixed with 4% paraformaldehyde, quenched with 0.1M glycine, permeabilized with 0.1% Triton X-100, and blocked with 10% donkey serum. Cells were incubated with chicken anti-GFP and AlexaFluor AF488-conjugated goat anti-chicken secondary antibodies (1:500), washed and mounted. At least 10 images of apical dendrites of EGFP-labeled pyramidal neurons were captured per condition. Confocal z-stacks were obtained using 0.2 µm optical sections of field size 64.02 × 64.02 µm with a 40x oil objective and 2.4x digital zoom, and subjected to deconvolution. Spines from maximum intensity projections were traced and scored blind to the observer using Neurolucida software. Mean spine densities (no./10 µm ± SEM) were calculated and compared by 2-factor ANOVA followed by Tukey’s posthoc pairwise comparisons with significance set at p < 0.05.

### Experimental Design and Statistical Analysis

All experiments were designed to provide sufficient power (80–90%) to discriminate significant differences (p<0.05) in means between independent controls and experimental subjects as described (40). Unpaired Mann-Whitney t-tests (2-tailed, unequal variance assumption) and ANOVA with Tukey’s post hoc multiple comparisons were calculated with Graphpad Prism. The type I error probability associated with these tests of the null hypothesis was set at 0.05.

## Results

### Increased Spine Density on Apical Dendrites of Cortical Pyramidal Neurons in L1 Null (L1-/y) Mutant Mice

To investigate a potential role for L1 in dendritic spine regulation we focused on pyramidal neurons in layer 2/3 of the medial prefrontal cortex (PFC, primary cingulate area) due to its importance in cognitive functions (41). In addition, two sensory cortical areas were studied: layer 5 of the primary motor (M1) and layer 4 of the visual cortex (V1), which are known to receive thalamic input. Brains of WT and hemizygous L1 null (L1-/y) male mice were sparsely labeled by Golgi-Cox impregnation in early adulthood (P50). Examination of Golgi-impregnated WT and L1-/y cerebral cortices showed typical pyramidal neurons in each cortical area with well-developed, branched apical dendrites that reached layer I, as well as basal dendrites extending from the soma. Only in M1, layer 5 of mutant mice were a minor number of apical dendrites laterally oriented as previously described (42), but these were not analyzed. Spine densities were quantified on the first branch of apical dendrites of pyramidal neurons using Neurolucida software. Spine densities were found to be significantly increased in PFC (layer 2/3), M1 (layer 5), and V1 (layer 4) in L1-/y mice compared to pyramidal neurons of WT mice in each cortical area (Mann-Whitney t-test, p < 0.05; Fig. 1 A-C). In contrast, the spine density on basal dendrites of L1-/y pyramidal neurons did not differ from WT in each of the areas (Fig.1 D-F). Apical dendrites are known to differ from basal dendrites in receiving different synaptic inputs and exhibiting distinct synaptic plasticity functions (43). Moreover, Neuropilin-2 is localized on apical but not basal dendrites of pyramidal neurons, accounting for Sema3-induced spine retraction specifically on apical dendrites (44).

**Figure 1:**
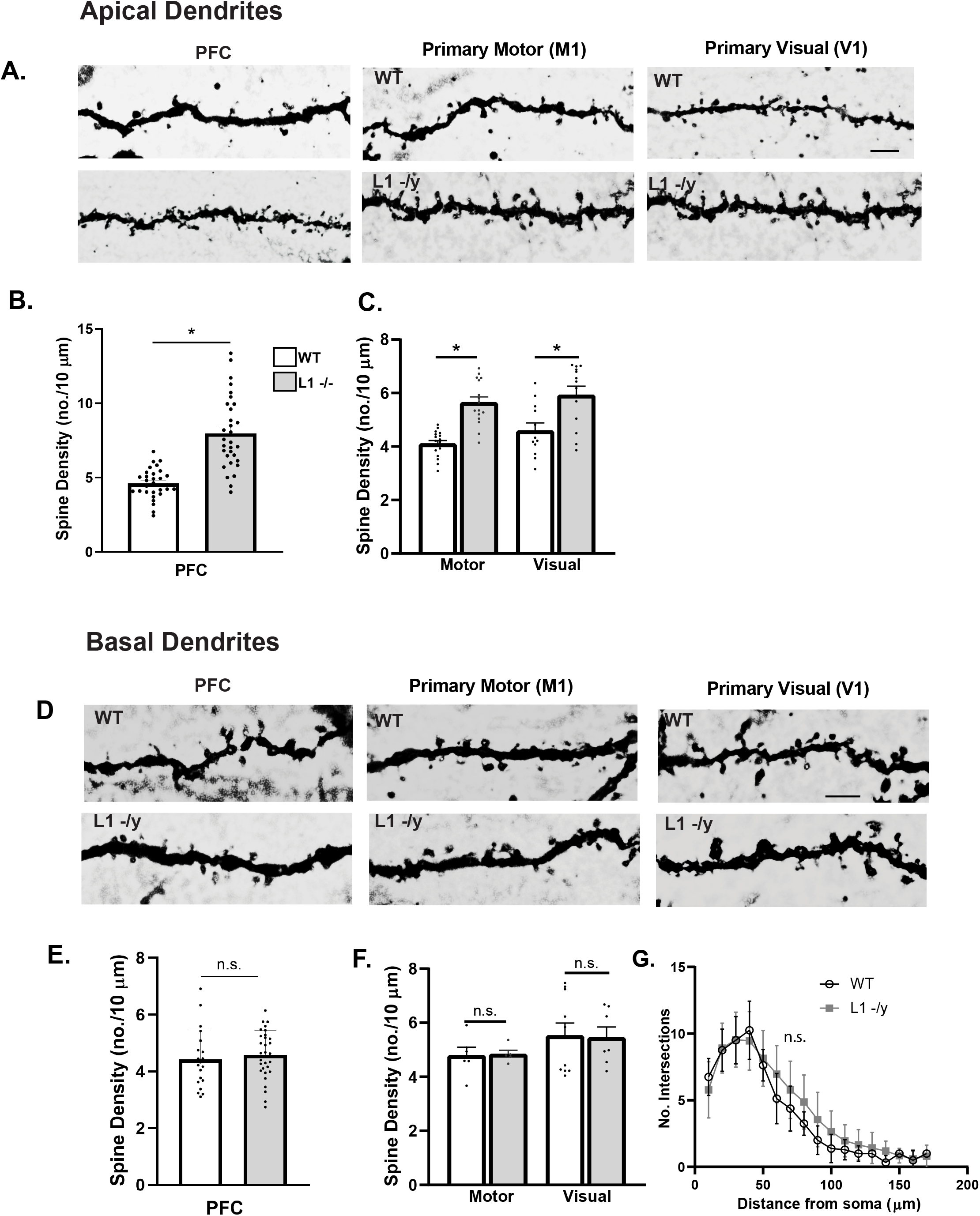
L1-/y null mutant mice display increased spine density on apical but not basal dendrites of cortical pyramidal neurons. (A) Representative images of apical dendrites of Golgi-labeled pyramidal neurons in PFC layer 2/3 (cingulate), primary motor cortex (M1, layer 5) and primary visual cortex (V1, layer 4) in WT or L1-/y mice (scale bar = 5 µm). (B, C) Mean spine densities of pyramidal neurons on apical dendrites of L1-/y PFC, M1, and V1 was significantly increased compared to WT (n ≥ 30 neurons/mouse; 6 mutant and 6 WT mice. PFC WT (4.6 ± 0.2 spines/10 μm); PFC L1-/y (8.0 ± 0.4); M1 WT (4.1/ 10 μm ± 0.1); M1 L1-/y (5.6 ± 0.2); WT V1 (4.6 ± 0.3); L1-/y V1 (5.9 ± 0.3). *PFC, p = 0.026; M1, p < 0.001; V1, p = 0.003 (Mann Whitney t-test). (D) Representative images of basal dendrites of Golgi-labeled pyramidal neurons in PFC layer 2/3; M1, layer 5; and V1, layer 4 of WT and L1-/y mice (scale bar = 5 µm). (E, F) Mean spine densities on basal dendrites of pyramidal neurons in L1-/y PFC, M1, and V1 were not significantly different (n.s., p > 0.05, Mann-Whitney t-test) compared to WT (n= 20-30 neurons; 6 mutant and 6 WT mice). WT PFC (4.4 ± 0.2 spines/10 μm); L1-/y PFC (4.6 ± 0.2); WT M1 (4.8 ± 0.3); L1-/y M1 (4.8 ± 0.1); WT V1 (5.5 ± 0.4); L1-/y V1 (4.6 ± 0.2). *PFC, p = 0.133; M1, p = 0.450; V1, p= 0.455 (Mann Whitney t-test). (G) There was no significant difference(n.s.) in arborization of dendrites in layer 2/3 pyramidal neurons in L1-/y PFC compared to WT shown by Sholl analysis (two factor ANOVA p = 0.12; n ≥ 10 neurons/mouse; 3 WT and 3 L1-/y mice).

To determine if loss of L1 affected dendritic arborization or spine morphology, we focused on pyramidal neurons in layer 2/3 of the PFC. Sholl analysis was performed on the dendritic tree of Golgi-labeled pyramidal neurons within the WT and L1-/y PFC. The dendritic arborization of mutant pyramidal neurons was not significantly different from WT in the PFC (layer 2/3) (Fig. 1 G, two factor ANOVA, p = 0.12). These results showed that L1 loss resulted in elevated spine density on apical rather than basal dendrites of pyramidal neurons in PFC (layer 2/3), M1 (layer 5), and V1 (layer 4). Dendritic protrusions acquire different morphologies and have been classified as mushroom, stubby, and thin spines based on the relative size of the spine head and neck (39). Spine morphologies are dynamically interchangeable, and comprise a continuum from thin spines, which can have excitable synapses with postsynaptic densities to mushroom spines with mature synaptic function (10, 45, 46). The density of spines with mushroom, stubby, or thin morphology on apical dendrites of Golgi-labeled pyramidal neurons in layer 2/3 of the PFC was compared between the WT and L1-/y genotypes at P50. L1-/y pyramidal neurons exhibited a significant increase in the density of mushroom spines (3.4 ± 0.4 spines/10 µm) compared to WT (2.4 ± 0.5 spines/10 µm) (Mann-Whitney t-test, p = 0.03). There was no significant difference in the density of stubby spines (0.4 ± 0.2 spines/10 µm) compared to WT (1.0 ± 0.1 spines/10 µm) (Mann-Whitney t-test, p = 0.97); or thin spines (0.5 ± 0.1 spines/10 µm) compared to WT (0.4 ± 0.09 spines/10 µm) (Mann-Whitney t-test, p = 0.53).

In summary we observed an increase in spine density on apical, not basal, dendrites of pyramidal neurons in all cortical areas of L1-/y mice compared to WT, and this was reflected in an increase in mature mushroom spines as shown in PFC layer 2/3.

### Increased Spine Density on Apical Dendrites of Cortical Pyramidal Neurons in L1 Y1229H Mutant Mice Mutated at the Ankyrin Binding Motif FIGQY

To determine if L1 interaction with Ankyrin was required for dendritic spine regulation we analyzed the PFC of L1Y1H knock-in mice, in which a histidine substitution for tyrosine at position 1229 causes deficiency in binding the actin-spectrin scaffold protein Ankyrin (28). Pyramidal neurons in WT and homozygous L1YH adult mice were sparsely labeled by Golgi-Cox impregnation and examined in layer 2/3 of PFC. WT and L1-/y pyramidal neurons displayed normal morphology and distribution (Fig. 2A). Spine densities were quantified on apical dendrites and found to be significantly increased in L1YH mice compared to WT (Fig. 2 B,C). However, spine density on basal dendrites of L1YH pyramidal neurons did not differ from WT (Fig. 2 B,D). To determine if mutation of the L1-Ankyrin binding motif affected dendritic arborization, Sholl analysis was performed on the dendritic tree of Golgi-labeled pyramidal neurons in layer 2/3 of WT and L1YH PFC. The dendritic arborization of mutant pyramidal neurons was not significantly different from WT (Fig. 2 E; two factor ANOVA, p = 0.95). To investigate whether the L1YH mutation affected spine morphology, the density of spines with mushroom, stubby, or thin morphology was quantified on apical dendrites of Golgi-labeled pyramidal neurons in PFC layer 2/3 of WT and L1YH mice. L1YH pyramidal neurons showed a significant increase in the density of mushroom spines (3.7 ± 0.7 spines/10 µm) compared to WT (3.1 ± 0.2) (Mann Whitney t-test, p=0.05). The density of stubby spines in L1YH mutants (1.3 ± 0.2 spines/10 µm) compared to WT (0.8 ± 0.09) (p = 0.40) was not significantly different, nor was the density of thin spines in L1YH mice (0.5 ± 0.08 spines/10 µm) compared to WT (0.6 ± 0.16) (p = 0.86).

**Figure 2:**
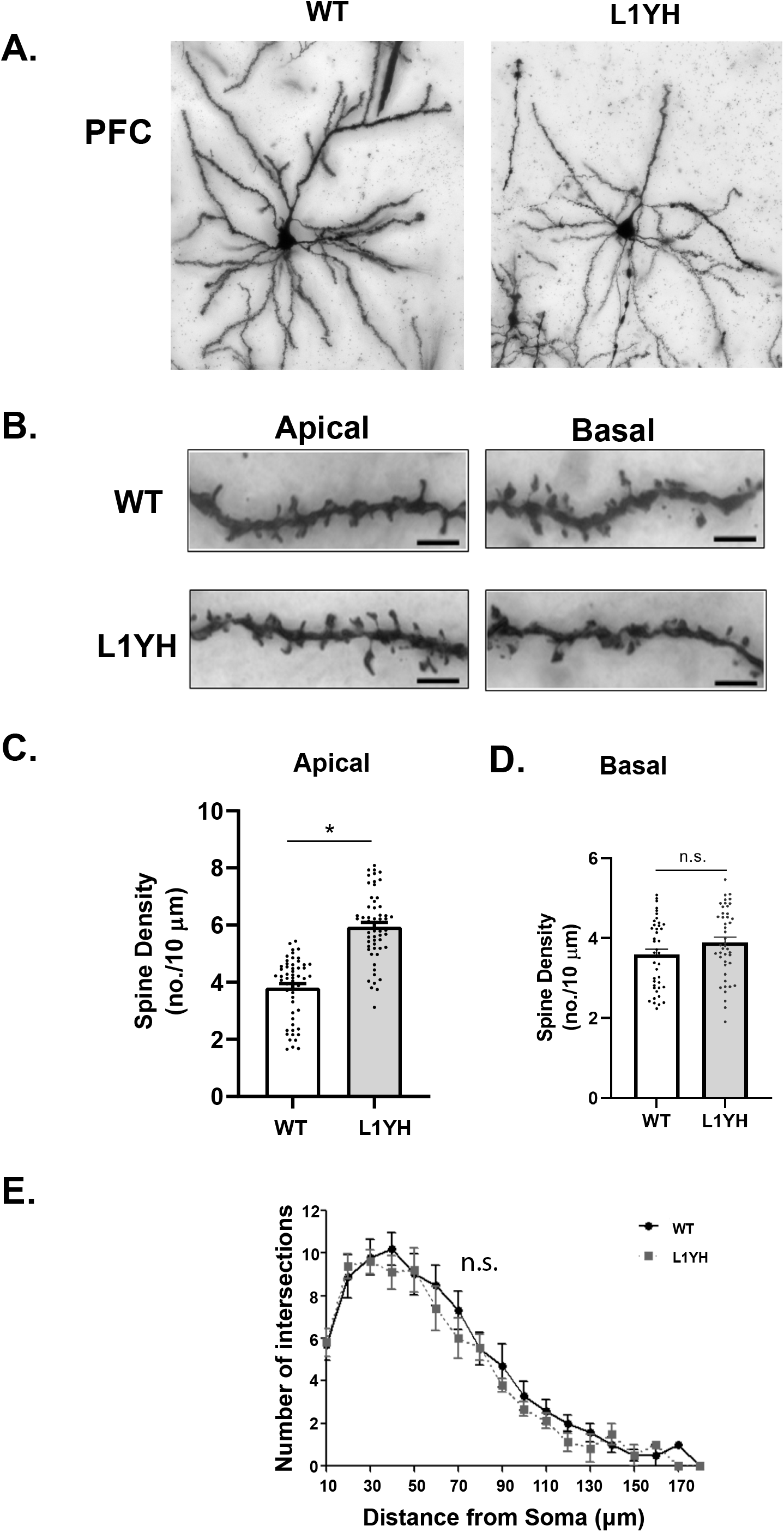
L1YH mutation increases spine density on apical dendrites but has no effect on dendritic branching. (A) Representative images of Golgi-labeled pyramidal neurons in PFC layer 2/3 of WT and L1YH mice. (B)Representative images of apical and basal dendrites of Golgi-labeled pyramidal neurons in PFC layer 2/3 of WT and L1YH mice (scale bar = 8 µm) (C)Mean spine densities on apical dendrites of layer 2/3 pyramidal neurons in the L1YH PFC were significantly increased compared to WT (n ≥ 55 neurons; n = 8 L1YH and 6 WT mice). WT (3.8 ± 0.1 spines/10 μm); L1YH (5.9 ± 0.2). *p < 0.001, Mann Whitney t-test. (D)Mean spine density of layer 2/3 pyramidal neurons in PFC on basal dendrites of L1YH was not significantly different compared to WT: WT (3.6 spines/10 μm ± 0.01), L1YH (3.9 ± 0.01). 8 L1YH and 6 WT mice.* p = 0.092, Mann Whitney t-test. (E)There was no significant difference (n.s.) in dendritic arborization of PFC layer 2/3 pyramidal neurons in L1YH compared to WT mice, as shown by Sholl analysis (two factor ANOVA p = 0.95; 3 WT and 3 WT mice, n ≥ 10 neurons/mouse).

In summary, we observed increased spine density on apical, not basal, dendrites of pyramidal neurons in PFC layer 2/3 of L1YH compared to WT mice, which was reflected in an increase in mushroom spines. Thus, both L1-/y and L1YH mice displayed increased spine densities primarily in mature spines on apical dendrites, with little or no effect on density of immature spines with stubby and thin morphology. These results suggest that L1 and its ability to recruit Ankyrin, play important roles in limiting the number of spines on apical dendrites of pyramidal neurons in PFC layer 2/3.

### L1 Expression in the PFC, Association with AnkyrinB, and Response to Semaphorins in Cultured Cerebral Cortical Neurons

In the visual cortex of the mouse, L1 has been detected on axons and apical dendrites of pyramidal neurons at embryonic and postnatal stages, declining in adulthood (42). Similarly, in cultures of neuron-induced human embryonic stem cells, L1 localized initially on all neurites and became restricted to axons upon further maturation (18). To determine if L1 was present in spines in the PFC *in vivo*, immunofluorescence staining of L1 was carried out in brain sections of WT mice (P20). WT transgenic mice (Nex1Cre: ERT2: RCE) were used because upon tamoxifen induction (P10-P13), they express EGFP in postmitotic, postmigratory pyramidal neurons, effectively labeling spines as well as dendrites, axons, and soma (32). L1 immunolabeling could be observed in spine heads in layer 2/3 pyramidal neurons of the PFC (Fig. 3A, three representative examples), as well as in dendrites identified by labeling with the somatodendritic marker MAP2. In addition L1 co-immunoprecipitated with the principal Ankyrin B isoform of 220 kDa (AnkB220) from lysates of WT mouse cortex but not from L1YH cortex (P30) (Fig. 3B). This result was in agreement with co-immunoprecipitation of L1 and AnkB220 from WT but not L1YH lysates of mouse superior colliculus (P8) (28). The L1YH mutation did not alter AnkB220 stability, as equal amounts of protein from WT and L1YH cortical lysates (inputs) showed equivalent levels of AnkB220 (Fig. 3B).

**Figure 3.**
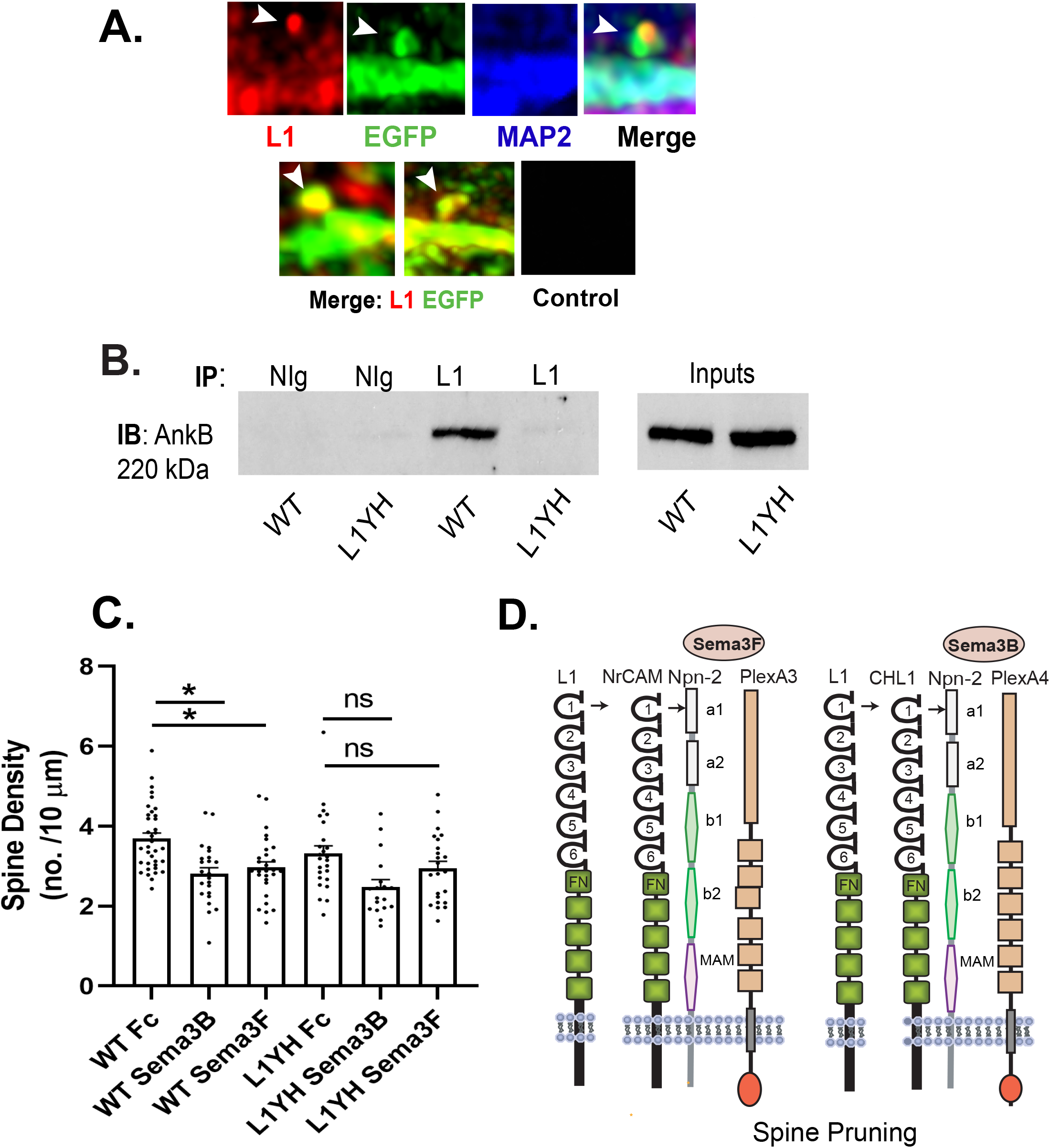
L1 expression and association with AnkyrinB in mouse neocortex, and spine retraction of WT and L1YH cortical neurons to Semaphorin 3B and 3F. (A) Immunofluorescence labeling of L1 (red) in spine heads (arrows) and dendrites (MAP2 immunolabeling, blue) of layer 2/3 pyramidal neurons in PFC of WT Nex1Cre-ERT2: RCE mice (P20) expressing EGFP (green). Two other representative merged images of L1 and EGFP immunofluorescence in spines are shown. Control is merged images of staining with all three secondary antibodies alone. (B) Co-immunoprecipitation of L1 and AnkyrinB (AnkB 220 kDa) with L1 antibodies from cortical lysates of WT but not L1YH mice (P30). IP, immunoprecipitation; IB, immunoblotting. NIg = nonimmune IgG. Inputs of cortical lysates (10 µg each) immunoblotted for AnkB 220 kDa showed equivalent amounts in WT and L1YH cortical lysates. (C)WT and L1YH cortical neuronal cultures were transfected with pCAG-IRES-EGFP, treated for 30 min with 5 nM Fc, Sema3B-Fc, or Sema3F-Fc on DIV14, and immunostained for EGFP. Apical dendrites were imaged confocally and spine density scored. Each point represents the mean spine density per 10 µm of dendrite on each neuron analyzed. Two factor ANOVA with Tukey’s posthoc test comparisons (* p< 0.05) showed that Sema3B-Fc and Sema3F-Fc significantly decreased spine density on WT but not L1YH cortical neurons compared to Fc control treated cultures (n.s, not significant). (WT Fc vs. Sema3B-Fc, *p=0.003; WT Fc vs. Sema3F-Fc, *p=0.011; L1YH Fc vs. Sema3B-Fc, p= 0.064; L1YH Fc vs. Sema3F-Fc, p=0.593). The data in the graph represents combined data from 3 experiments with different neuronal cultures. (D)Scheme showing postulated functional redundancy of L1 with NrCAM or CHL1 for binding to Neuropilin2 (Npn-2) in Semaphorin3 receptor complexes in the spine plasma membrane. Sema3F-induced spine pruning is mediated by the holoreceptor comprising NrCAM, Npn-2, and PlexinA3 (PlexA3), while the response to Sema3B is mediated by the holoreceptor comprising CHL1, Npn-2, and PlexinA4 (PlexA4). Immunoglobulin (Ig)-like domains 1-6 and fibronectin III domains (FN) on L1 family members, and Npn-2 domains a1, a2, b1, b2, and MAM domains are shown.

To investigate if the L1-Ankyrin interaction was required for Sema3F-or Sema3B-mediated spine retraction in an *in vitro* system, cortical neuron cultures were prepared from forebrains of WT and L1YH embryos (E15.5) and cultured for 14 days *in vitro* (DIV) as described (33). On DIV11 cells were transfected with pCAG-IRES-EGFP to enhance spine visualization, and treated on DIV14 with Sema3F-Fc, Sema3B-Fc or control Fc proteins (5nM) for 30 minutes (32, 34). Spine densities on EGFP-labeled apical dendrites were quantified and mean spine densities compared. Several reports have documented that Sema3B-Fc and Sema3F-Fc promote spine retraction in WT cortical neurons in culture (32-34). As expected Sema3B-Fc and Sema3F-Fc induced spine retraction, decreasing the mean spine density on apical dendrites in WT cultures to a significant extent (Fig. 3C). Spine density was not significantly different between WT and L1YH control cultures (Fc treated, p = 0.633). In L1YH cultures, Sema3B-Fc appeared to induce spine retraction compared to Fc-treated L1YH neurons but this difference was not statistically significant (p = 0.064). Sema3F-Fc did not significantly induce spine retraction in L1YH neurons compared to Fc-treated L1YH neurons (p = 0.593).

Spine morphology of WT and L1YH cortical neurons was quantified in Fc-treated cultures to determine if mutation of the L1 Ankyrin binding motif altered the percent of mushroom, stubby or thin spines) *in vitro*. There were no significant differences in the percent of mushroom, stubby, or thin spines in WT (33%, 25%, 42%, respectively) compared to L1YH cultures (35%, 25%, 39%) (2 tailed t-test, unequal variance, p< 0.05).

In summary, the results of the neuronal culture assays suggested that Sema3F, and possibly Sema3B, promote spine retraction through L1 association with Ankyrin. It should be noted that L1-/y neuronal cultures were not analyzed, because breeding requires mating WT males with heterozygous L1-/+ females which produces litters with a low percentage of L1-/y male embryos.

## Discussion

Using the L1-null mouse model we show that the neural cell adhesion molecule L1 constrains dendritic spine density in pyramidal neurons in the mouse cerebral cortex. This novel function for L1 was restricted to apical dendrites of pyramidal neurons and preferentially targeted mature mushroom spines. The Ankyrin binding site in the L1 cytoplasmic domain was required for constraining spine density as demonstrated by increased spine density in the mouse mutant containing the L1 YH mutation. This tyrosine to histidine substitution in the L1 cytoplasmic domain perturbs L1-Ankyrin binding and is a known variant associated with the human L1 syndrome of intellectual disability (19).

The present study extends the function of L1 to dendritic spine regulation from its well-established role in axon growth and guidance (14, 15). The increased density of spines in L1-/y and L1YH mice on apical and not basal dendrites in diverse cortical areas (PFC, M1, V1) indicates that spine regulation is impacted by L1 deficiency or loss of L1 binding to Ankyrin. These findings indicated that L1 and its interaction with Ankyrin likely promotes spine pruning, although inhibition of spine formation cannot be excluded. Mature mushroom spines were increased in the two L1 mutant mouse models suggesting that they may be preferentially targeted. *In vitro* assays showed that the L1-Ankyrin interaction supported spine retraction to Sema3F-Fc, and a nearly significant trend for Sema3B-Fc. As shown in Fig. 3D, NrCAM binds Neuropilin2 (Neuropilin1 not tested) at a site (TARNER) in its immunoglobulin (Ig) – like Ig1 domain necessary for Sema3F-induced spine retraction (32). CHL1 binds Neuropilin2 (and Neuropilin1) at a homologous sequence (FASNKL) in its Ig1 domain (47) for Sema3B-induced spine pruning (34). As depicted in Fig. 3D, L1 may engage Neuropilin2 to promote spine pruning in response to Sema3F and Sema3B, as the L1 Ig1 domain contains a homologous sequence (FASNKL) that binds Neuropilins, which is necessary for Sema3A-induced growth cone collapse (48, 49). This may result in functional compensation or competition among L1-CAMs for spine pruning. Similarly, functional redundancy among L1 family members has been demonstrated for L1 and CHL1 in thalamocortical axon guidance and topographic mapping (50). In regard to other Semaphorins, Sema3A does not induce spine retraction in cortical neuron cultures (34), and mice deficient in Sema3A exhibit normal spine density (44). Sema3C, Sema3D, and Sema3E also had no effect on spine density *in vitro* (34).

The spine phenotype of L1YH mice reveals a novel role for L1-Ankyrin binding in constraining spine density in pyramidal neurons in the PFC. Previously, L1YH mutants were shown to display errors in retinocollicular axon guidance and synaptic targeting (28). Ankyrin may position L1 on the cell surface to respond to Semaphorins or other factors regulating spine remodeling. L1-Ankyrin interaction also facilitate synapse formation or maintenance. Ankyrin binding to the L1 FIGQY site stabilizes synaptic connections between GABAergic interneurons and pyramidal cell soma (29) or axon initial segments (30). L1 is also involved in stabilizing perisomatic synapses of hippocampal neurons (51), and formation of inhibitory and excitatory synapses on cerebellar Purkinje cells (52).

A limitation of our study on L1-/y and L1 YH adult mice is that earlier stages at which L1 may act in spine regulation were not examined. L1 is known to be expressed at highest levels at postnatal stages in the mouse cortex and to decline with maturation (42), suggesting that L1 could function at earlier postnatal stages during the most active phase of spine remodeling. On the other hand, L1 may regulate adult spine plasticity, since mature mushroom spines preferentially increased in L1-/y and L1YH mice. Although we did not evaluate excitatory synapses or neurotransmission, CAMKII-Cre conditional mice targeting L1 in pyramidal neurons displayed increased basal excitatory transmission in the hippocampus (53). Increased excitatory connectivity could contribute to observed behavioral deficits in L1 mutant mice, which include decreased anxiety (53), altered sociability and increased repetitive behaviors (54).

In conclusion, the present study extends the role of L1 family members in dendritic spine regulation to the prototype of the family, L1, and its interaction with Ankyrin. The increased spine density due to L1 deficiency or loss of Ankyrin binding may alter excitatory/inhibitory balance in cortical circuits and affect overall behavior. The phenotypes observed in the L1 mouse genetic models studied here may also shed light on the molecular basis of cognitive and other L1-related functions that are abnormal in the L1 syndrome.

## Data Availability Statement

Mice and data supporting the findings of this work can be obtained from the authors.

### Author Contributions

KEM, SDW, JS, VM, BWD, and YP propagated L1YH mouse strains, performed experiments, and analyzed data. DL and MS supervised breeding of L1-/y mice, performed Golgi staining on brain sections of L1-/y mice, and provided advice on experimental design. PM supervised research, analyzed data, and wrote the paper. **Acknowledgements**

This work was supported by the US National Institutes of Mental Health grant R01 MH113280 (PFM), UNC School of Medicine Biomedical Research Core Project award (PFM), Carolina Institute for Developmental Disabilities center grant NIH P50HD103573 (Dr. Joseph Piven, PI), NIH T32 NRSA (5T32HD040127-18)(KEM), FoRUM grant F957N-2019 of the Ruhr University Bochum, Germany and the Germany DAAD grant (#5756078) (DL) and Deutsche Forschungsgemeinschaft (SCHA 185/87-3)(MS). We acknowledge Dr. Pablo Ariel, Director of the Microscopy Services Laboratory in the UNC Department of Pathology and Laboratory Medicine, who provided expert advice on imaging (P30 CA016086 Cancer Center Core Grant). We thank Alexander Kampov-Polevoi, Teva Smith, and Cassandra Sweetman for assistance with experiments. Gabriele Loers and Eva Kronberg are kindly acknowledged for L1-/y mouse breeding.

